# C2CD6 is required for assembly of the CatSper calcium channel complex and fertilization

**DOI:** 10.1101/2021.07.06.451342

**Authors:** Fang Yang, Maria Gracia Gervasi, N. Adrian Leu, Darya A. Tourzani, Gordon Ruthel, Pablo E. Visconti, P. Jeremy Wang

## Abstract

The CatSper cation channel is essential for sperm capacitation and male fertility. The multi-subunit CatSper complexes form highly organized calcium signaling nanodomains on flagellar membranes. Here we report identification of an uncharacterized protein C2CD6 as a novel subunit of the CatSper ion channel complex. C2CD6 contains a calcium-dependent membrane targeting C2 domain. C2CD6 interacts with the CatSper calcium-selective core forming subunits. Deficiency of C2CD6 depletes the CatSper nanodomains from the flagellum and results in male sterility. C2CD6-deficient sperm are defective in hyperactivation and fail to fertilize oocytes both in vitro and in vivo. Interestingly, transient treatments with either Ca^2+^ ionophore, starvation, or a combination of both restore the fertilization capacity of C2CD6-deficient sperm in vitro. C2CD6 interacts with EFCAB9, a pH-dependent calcium sensor in the CatSper complex. We postulate that C2CD6 may regulate CatSper assembly, target the CatSper complex to flagellar plasma membrane, and function as a calcium sensor. The identification of C2CD6 as an essential subunit may facilitate the long-sought reconstitution of the CatSper ion channel complex in a heterologous system for male contraceptive development.

## Introduction

Sperm acquires fertilization competence in the female reproductive tract (Chang, 1951). Sperm hyperactivated motility triggered by capacitation is essential for navigation in the oviduct (Suarez, 2016), rheotaxis (Miki and Clapham, 2013), and zona pellucida penetration (Stauss et al., 1995). The uterus and oviduct fluids provide an alkaline environment and a high concentration of bicarbonate, which are critical for capacitation (Vishwakarma, 1962). Sperm motility is regulated by ion channels and ion transporters in the flagellum in response to environmental stimuli (Vyklicka and Lishko, 2020). Calcium influx in sperm is gated by the CatSper ion channel in the flagellum (Ren et al., 2001). The CatSper channel is activated by alkaline pH in rodents (Kirichok et al., 2006) and primates (Lishko et al., 2010), and by progesterone (P4) in primates (Lishko et al., 2011;Strunker et al., 2011). The CatSper channel forms four linear columns of Ca^2+^ signaling domains along the principal piece of the sperm flagellum (Chung et al., 2014). The CatSper domains organize the spatiotemporal pattern of tyrosine phosphorylation of flagellar proteins, one of the hallmarks of capacitation (Chung et al., 2014;Visconti, P. E. et al., 1995). Calcium influx caused by the CatSper activation results in powerful asymmetrical flagellar beating movement known as hyperactivation. CatSper is inhibited by efflux of potassium, which is carried out by the Slo3 K^+^ channel (Brenker et al., 2014;Chavez et al., 2014;Geng et al., 2017;Santi et al., 2010;Schreiber et al., 1998;Zeng et al., 2011). In human spermatozoa, P4 binds to its sperm membrane receptor ABHD2, which hydrolyzes the endocannabinoid 2-arachidonoylglycerol (2-AG), an inhibitor of the CatSper channel. As a result, P4 activates CatSper by removing 2-AG from the plasma membrane (Miller et al., 2016). Therefore, sperm hyperactivation is regulated by both environmental stimuli in the female reproductive tract and ion channels on sperm flagella.

The CatSper channel is a complex of ten known subunits: CatSper1-4, CatSperβ, γ, δ, ε, ζ, and EFCAB9 (Lin et al., 2021;Vyklicka and Lishko, 2020;Wang et al., 2021). CatSper1-4 subunits form a heteromeric complex with a central Ca^2+^-selective pore. The remaining six subunits are auxiliary proteins. While CatSperβ, γ, δ, and ε are putative transmembrane proteins, CatSperζ and EFCAB9 lack transmembrane domains (Chung et al., 2011;Chung et al., 2017;Hwang et al., 2019;Liu et al., 2007;Wang et al., 2009). Genetic ablation in mice and humans have revealed the role of the CatSper subunits in male fertility. Each of the CatSper core subunits (CatSper1-4) is required for the CatSper complex formation, sperm hyperactivation, and thus, for male fertility (Carlson et al., 2003;Carlson et al., 2005;Jin et al., 2007;Qi et al., 2007;Quill et al., 2003;Ren et al., 2001). Like CatSper1-4, CatSperδ, is essential for CatSper channel complex assembly and male fertility (Chung et al., 2011). In contrast, CatSperζ or EFCAB9-deficient males exhibit subfertility (Chung et al., 2017;Hwang et al., 2019). In CatSperζ or EFCAB9-deficient mouse mutants, the CatSper Ca^2+^ signaling domain organization is affected but the CatSper channel is still functional. EFCAB9 is an EF-hand calcium binding protein. EFCAB9 interacts with CatSperζ and this interaction requires the binding of Ca^2+^ to EF-hand domains. Thus, EFCAB9 in partner with CatSperζ functions as an intracellular pH dependent Ca^2+^ sensor and activator for the CatSper channel (Hwang et al., 2019). Each of the four columns of CatSper domains consists of two rows. However, in CatSperζ or EFCAB9-deficient mouse sperm, each column contains only one row of CatSper domains instead of two, suggesting a structural role in addition to their Ca^2+^ sensor function. The CatSper channel is essential for male fertility in humans. Men with loss of function mutations in CatSper subunits are infertile due to failures in sperm hyperactivation (Avenarius et al., 2009;Brown et al., 2018;Luo et al., 2019;Smith et al., 2013).

CatSper is probably the most complex ion channel known to date. Despite the extensive genetic, super-resolution structural, and electrophysiological studies, a functional CatSper complex has not been reconstituted in a heterologous system. One possibility is that additional subunits are yet to be identified. Serendipitously, we identified a calcium-binding C2 membrane domain protein (C2CD6) as a novel subunit for the CatSper channel complex. Here, we demonstrate that C2CD6, like CatSper1-4 and CatSperδ, is essential for the CatSper assembly, sperm hyperactivation, and male fertility.

## Materials and Methods

### Generation of *C2cd6* knockout mice

The targeting strategy was to delete 1.6-kb genomic region including exon 1 of the *C2cd6* gene (Fig. 2A). In the targeting construct, the left (2.3 kb) and right (2.1 kb) homologous arms were PCR amplified from a *C2cd6*-containing mouse BAC clone (RP24-535E21) with high-fidelity Taq DNA polymerase. A neomycin (PGKNeo) selection marker was inserted between the homologous arms. The HyTK selection marker was cloned adjacent to the right arm. V6.5 embryonic stem (ES) cells were electroporated with the ClaI-linearized targeting construct and cultured in the presence of 350 μg/ml G418 and 2 μM ganciclovir. ES cell clones were screened by long-distance PCR for homologous recombination. Out of 192 ES cell clones, 9 homologously targeted clones were identified. 2C5 and 1H1 ES cell clones were injected into blastocysts and the resulting chimeric mice transmitted the *C2cd6* knockout allele through the germline. The *C2cd6*^+/-^ 2C5 mice were backcrossed to the C57BL/6J strain four times (N4). All the experiments were performed on the C57BL/6J N4 backcrossed mice. Mice were genotyped by PCR of tail genomic DNA with the following primers: wild type (515 bp), ALS-25 (5′-GTATTTCCCATCATGTGGAGGA-3′) and ALS-26 (5′-AGTGGCTTGCCTTCTTCATCAG-3′); *C2cd6* knockout allele (341 bp), ALS-9 (5′-TGTGCTATCCACCTTGCCTT-3′) and PGKRNrev2 (5′-CCTACCGGTGGATGTGGAATGTGTG-3′).

*CatSper1* knockout mice were previously generated (Ren et al., 2001). All experiments with mice were performed in accordance with the Institutional Animal Care and Use Committee (IACUC) guidelines of the University of Pennsylvania and the University of Massachusetts at Amherst.

### Antibody production

The *C2cd6* cDNA was amplified from bulk mouse testis cDNA by PCR. The cDNA fragment encoding residues 1-386 was cloned into the pQE-30 vector (QIAGEN). The 6×His-C2CD6 (aa 1-386) fusion protein was expressed in M15 bacteria, affinity purified with Ni-NTA beads, and eluted in 8 M urea. The recombinant fusion protein was used to immunize two rabbits at Cocalico Biologicals, Inc. The C2CD6 antiserum (UP2429 and UP2430) was used for Western blotting analysis (1:500). Specific antibodies for immunofluorescence were affinity purified with the immunoblot method (Harlow and Lane, 1998).

The *Als2cr11b* cDNA fragment encoding the C-terminal 200 residues was cloned into the pQE-30 vector. The 6× His-ALS2CR11B (C-terminal 200 aa) fusion protein was expressed in bacteria, affinity purified, and used to immunize two rabbits at Cocalico Biologicals, In, resulting in one anti-ALS2CR11B antiserum (UP2443).

### In vivo fertilization

Eight-week-old wild type C57BL/6 females were injected with 7.5 IU of PMSG, then with 7.5 IU of hCG 46 hours later, and mated with either *C2cd6*^+/-^ or *C2cd6*^-/-^ males. Copulatory plugs were checked 17 hours after mating setup. 24 hours after plug check, eggs/embryos were flushed from plugged females. The numbers of two-cell embryos and one-cell embryos/eggs were counted.

### Sperm collection

Cauda epididymides from three month-old males were dissected and placed in Toyoda-Yokoyama-Hosi (TYH) buffer containing 119.37 mM NaCl, 4.7 mM KCl, 1.71 mM CaCl_2_, 1.2 mM KH_2_PO_4_, 1.2 mM MgSO_4_, 25.1 mM NaHCO_3_, 0.51 mM sodium pyruvate, 5.56 mM glucose, 4 mg/ml BSA, 10 µg/ml gentamicin and 0.0006% phenol red equilibrated in 5% CO_2_ at 37°C. After 10 minutes of sperm swim-out, epididymal tissue was removed and sperm suspensions were further incubated in TYH in 5% CO_2_ at 37 °C to allow sperm to capacitate.

### Sperm motility analysis

Sperm motility parameters were analyzed immediately after swim-out (T0) and after 60 minutes (T60) of incubation in capacitating conditions (TYH medium). Sperm suspensions were loaded onto 100 µm depth chamber slides (Leja, Spectrum Technologies, CA), placed in a slide warmer at 37 °C (Minitherm, Hamilton Thorne), and imaged with a 4× dark field objective (Olympus) in a Lab A1 microscope (Zeiss). Ninety frame videos were recorded at 60 Hz and analyzed using a CEROS II computer assisted sperm analysis system (Hamilton-Thorne Inc., Beverly, MA). The settings for cell recognition included: head size, 5-200; min head brightness, 100; and static head elongation, 5–100 %. Sperm with average path velocity (VAP) > 0 and rectilinear velocity (VSL) > 0 were considered motile. Sperm were considered progressive with VAP > 50 µm/s and straightness (STR) > 50 %, and hyperactive with curvilinear velocity (VCL) > 271 µm/s, VSL/VCL (LIN) < 50 %, and amplitude of lateral head displacement (ALH) > 3.5 µm. At least five fields per treatment corresponding to a minimum of 200 sperm were analyzed per experiment. Data were presented as percentage of motile sperm out of the total population, and percentage of progressive or hyperactive sperm out of the motile population.

### In vitro fertilization (IVF)

Young (7-9 weeks-old) CD-1 females were obtained from Charles River Laboratories (Wilmington, MA). Superovulation was induced by injection first with 7.5 IU of PMSG (Cat. No. 493-10, Lee Biosolution), followed 48 hours later with 7.5 IU of hCG (Cat. No. CG5, Sigma). Females were sacrificed 13 hours post-hCG injection, the oviducts were dissected, and cumulus-oocyte complexes (COCs) were collected in TL-HEPES medium containing 114 mM NaCl, 3.22 mM KCl, 2.04 mM CaCl_2_, 0.35 mM NaH_2_PO_4_, 0.49 mM MgCl_2_, 2.02 mM NaHCO_3_, 10 mM lactic acid (sodium salt), and 10.1 mM HEPES. COCs were thoroughly washed with TYH and placed in an insemination drop of TYH covered by mineral oil previously equilibrated in an incubator in 5% CO_2_ at 37 °C. COCs (2-3 per 90 μl drop) were inseminated with 100,000 sperm that were previously capacitated in TYH medium for 60 minutes, and then were maintained in an incubator in 5% CO_2_ at 37°C. After 4 hours of insemination, the MII-oocytes were washed, placed in a different drop of TYH media, and incubated overnight in 5% CO_2_ at 37 °C. Dishes were examined 24 hours post-insemination and fertilization was evaluated by the appearance of two-cell stage embryos. Results were expressed as percentage of 2-cell embryos out of the total number of oocytes inseminated.

### Enhanced IVF

Sperm Energy restriction and Recovery (SER) and Ca^2+^ ionophore treatments prior to IVF improve the rate of fertilization in subfertile and infertile animals (Navarrete et al., 2016;Navarrete et al., 2019). The following sperm treatments prior to IVF were tested: 1) Control (TYH); 2) Control + Ca^2+^ ionophore; 3) SER; and 4) SER + Ca^2+^ ionophore. For each 3-month-old male, one cauda epididymis was collected and placed in TYH medium (tube A) and the other cauda epididymis was collected and placed in TYH medium devoid of glucose and pyruvate (SER-TYH, tube B). After 10 minutes of incubation in 5% CO_2_ at 37 °C to allow sperm swim-out, cauda tissues were removed. Sperm suspensions were centrifuged twice at 150 g for 5 minutes and the sperm pellet was washed with 2 ml of TYH for tube A and 2 ml of SER-TYH for tube B. After the final wash, sperm pellets were resuspended in 500 µl of TYH for tube A and 500 µl of SER-TYH for tube B. Each tube was immediately divided into two 250 µl suspensions and incubated in 5% CO_2_ at 37 °C until the sperm motility from treatments 3 and 4 was significantly slow (about 30 minutes). At that point, Ca^2+^ ionophore 4Br-A23187 (Cat. No. C7522, Fisher Scientific) was added to a final concentration of 20 µM for treatment 2 and 5 µM for treatment 4. After 10-minute incubation with Ca^2+^ ionophore, 1.5 ml of TYH was added to all the treatments and immediately centrifuged at 150 g for 5 min. Sperm pellets were then washed again with TYH and centrifuged at 150 g for 5 minutes. Sperm pellets were resuspended in 500 ul of TYH and used for insemination of COCs. All the procedures after insemination were performed as described above for regular IVF.

### Embryo culture

The two-cell embryos from IVF were washed and placed in a dish of equilibrated KSOM media (Cat. No. MR-106-D, Fisher Scientific) covered by light mineral oil (Cat. No. 0121-1, Fisher Scientific). Embryo culture dishes were incubated for 3.5 days in 5% CO_2_, 5% O_2_, at 37°C. Results are expressed as percentage of blastocysts out of the 2-cell embryos and percentage of blastocysts out of the total number of oocytes inseminated.

### Immunofluorescence and super-resolution imaging

After three incisions, cauda epididymides were incubated in PBS at 37°C for 10 min. Swim-out sperm were placed on slides and fixed in 2% PFA in PBS with 0.2% Triton X-100 overnight. The slides were incubated with anti-C2CD6 (UP2430, 1:100) or anti-Catsper1 (1:200) antibodies, then with anti-rabbit FITC-conjugated secondary antibody, and finally mounted with DAPI. Images were captured with an ORCA digital camera (Hamamatsu Photonics) on a Leica DM5500B microscope. For structured illumination microscopy (SIM) imaging, images were acquired on a GE DeltaVision OMX SR imaging system with PCO sCMOS cameras and were processed using softWoRx software.

### Testis microsome extraction

One adult testis (∼100 mg) was homogenized on ice in 1ml 0.32 M sucrose solution with 1× protease inhibitor cocktail (Cat No. P8340, Sigma). After centrifugation at 300 g at 4°C for 10 minutes, the supernatant was transferred to an ultra-centrifuge tube and centrifuged at 100,000 g for one hour. The pellet containing the microsome fraction was resuspended and solubilized in 5 ml PBS with 1% Triton X-100 and 1× protease inhibitor cocktail by rocking at 4°C for 2 hours. The suspension was centrifuged at 15,000 g for 30 minutes and the supernatant was collected for Western blot analysis.

### Sperm protein extraction

Sperm (∼1.3 × 10^7^) were collected from one adult mouse by squeezing the cauda epididymides in PBS solution and centrifugation at 800 g for 5 minutes at room temperature. Sperm were homogenized in 100 µl SDS-EDTA solution (1% SDS, 75 mM NaCl, 24 mM EDTA, pH 6.0) and centrifuged at 5000 g for 30 minutes at room temperature. 100 µl 2× SDS-PAGE sample buffer (62.5 mM Tris, pH 6.8, 3% SDS, 10% glycerol, 5% β-mercaptoethanol, 0.02% bromophenol blue) was added to 100 µl of supernatant. The samples were heated at 95°C for 10 minutes and 20 µl of each sample (equivalent to 1.3 ×10^6^ sperm) was used for Western blot analysis.

### Cell culture, transfection, and immunoprecipitation

The ORFs of *C2cd6s, Efcab9, CatSperζ, CatSper2, CatSper3*, and *CatSper4* were PCR amplified from bulk mouse testis cDNAs. The *CatSper1* ORF was amplified from a mouse cDNA clone (Cat. No. MR224271, Origene). *C2cd6s* was subcloned into pcDNA3.1/myc-His A vector (Cat. No. V800-20, Invitrogen). The others were TA-cloned to pcDNA3.1/V5-His TOPO TA vectors (Cat. No. K4800-01, Invitrogen). HEK293T cells were cultured in DMEM medium with 10% FBS in 5% CO_2_ at 37°C. 24 to 48 hours after transfection, the cells (3 wells of a 6-well plate) were lysed in 1 ml IP buffer (50 mM Tris, pH 8.0, 150 mM NaCl, 5 mM MgCl_2_, 1m M DTT, 0.5% deoxycholate, 1% Triton) with 1× cocktail of protease inhibitors, incubated at 4°C for one hour, and centrifuged at 12,000 g for 30 minutes. 10 µl of supernatant (1%) was set aside as input. The bulk of supernatant (∼1 ml) was incubated with antibodies at 4°C for one hour: 3 µl (0.5 µg/µl) c-Myc monoclonal antibody (Cat. No. 631206, TaKaRa), or 1 µl (1.1 µg/µl) anti-V5 antibody (Cat. No. P/N 46-0705, Invitrogen), or 20 µl anti-C2CD6 (UP2429) antibody. 10 µl of Dynabeads G or A (Invitrogen) was added for each IP and incubated at 4°C overnight. Immunoprecipitated proteins were washed five times with the wash buffer (50 mM Tris, pH 7.5, 250 mM NaCl, 0.1% NP-40, 0.05% deoxycholate) and were eluted in 15 µl 2× SDS sample buffer at 95°C. Elutes and inputs were separated by SDS-PAGE and transferred to nitrocellulose membrane. Anti-c-Myc monoclonal, anti-V5, and anti-CatSper1 (gift from Dejian Ren) (Ren et al., 2001) antibodies were used for Western blotting.

## Results

### Identification of a novel evolutionarily conserved sperm flagellar protein

We were interested in interacting proteins of TEX11, a meiosis-specific protein that we previously identified (Yang et al., 2008;Yang et al., 2015). ALS2CR11 was reported as one of the TEX11-interacting proteins in the genome-wide protein-protein interaction study (Rual et al., 2005). However, we find that ALS2CR11 (NP_780409) localizes to sperm flagellum but does not function in meiosis. Since ALS2CR11 contains a calcium-dependent membrane targeting C2 domain, it has been renamed as C2CD6 (<u>C2</u> <u>c</u>alcium dependent <u>d</u>omain containing <u>6</u>) (Fig. 1A and 1B) (Nalefski and Falke, 1996). The *C2cd6* gene is conserved in vertebrates. The transcripts from the *C2cd6* gene locus are very complex. As we were analyzing the *C2cd6* gene structure on Chromosome 1, we noticed another uncharacterized gene 3’ downstream, which has a large coding exon (∼5 kb), and named it *Als2cr11b* (Fig. 1A). Based on EST (expressed sequence tag) profiling, both *C2cd6* and *Als2cr11b* transcripts are testis-specific.

**Figure 1.**
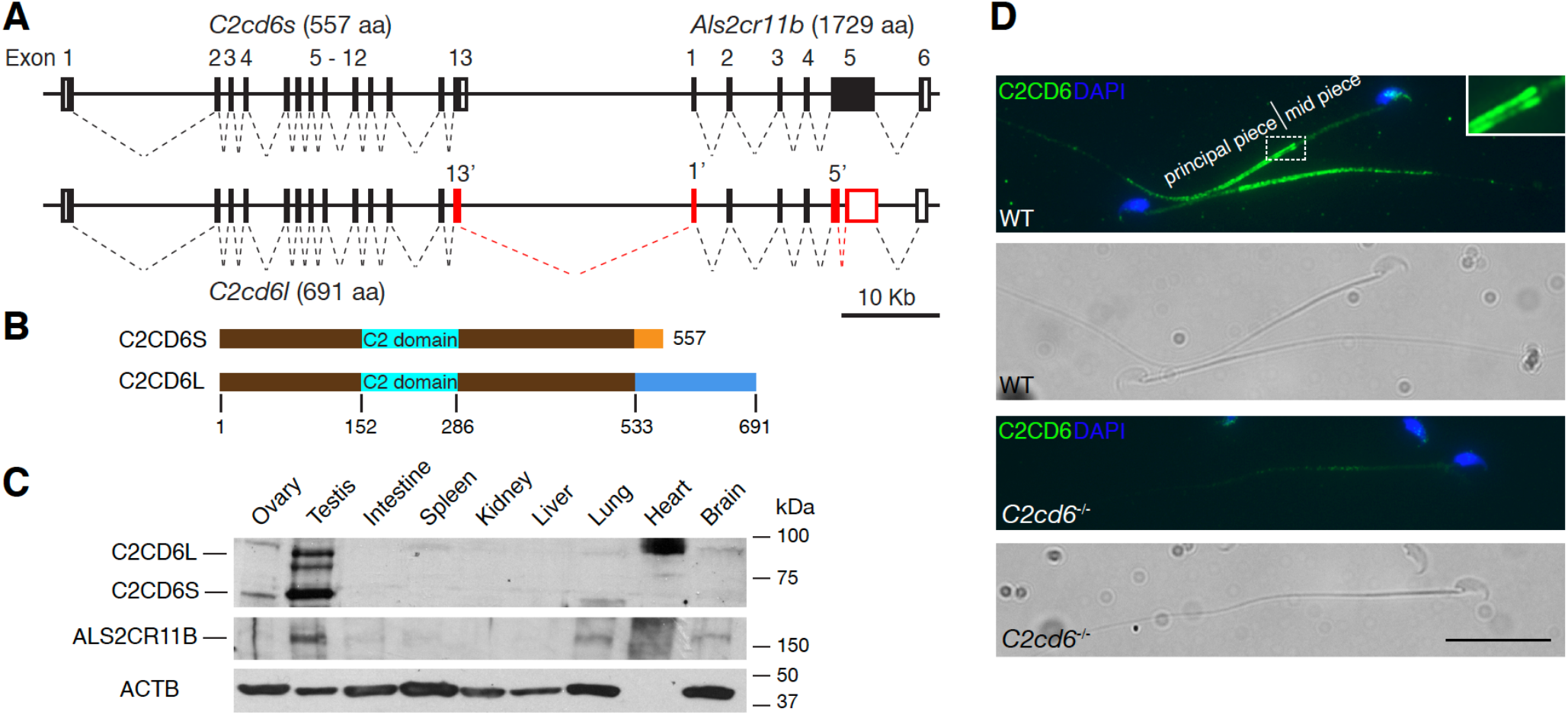
C2CD6 localizes to the principal piece of sperm flagella. (A) Gene structures of *C2cd6* and *Als2cr11b* on Chromosome 1. Coding regions of exons are shown in black boxes. 5′ and 3′ untranslated regions (UTR) are shown as open boxes. Alternative exons of *C2cd6l* are shown in red. Alternative exons 13′ and 1′ are in frame. Splicing of an intron within *Als2cr11b* exon 5 resulting in early frame shift in the *C2cd6l* transcript. GenBank accession numbers for cDNA sequences: *C2cd6l*, MW717645; *C2cd6s*, NM_175200; *Als2cr11b*, XM_011238634. (B) Schematic diagram of two C2CD6 protein isoforms. The N-terminal 533 residues are identical. The Ca^2+^ binding membrane targeting C2 domain is predicted based on Phyre2. (C) Western blot analysis of C2CD6 and ALS2CR11B in adult mouse tissues. ACTB serves as a loading control. Note that heart lacks ACTB. (D) Localization of C2CD6 to the principal piece of wild type mouse sperm but not the *C2cd6*-null sperm. Scale bar, 25 μm.

**Figure 2.**
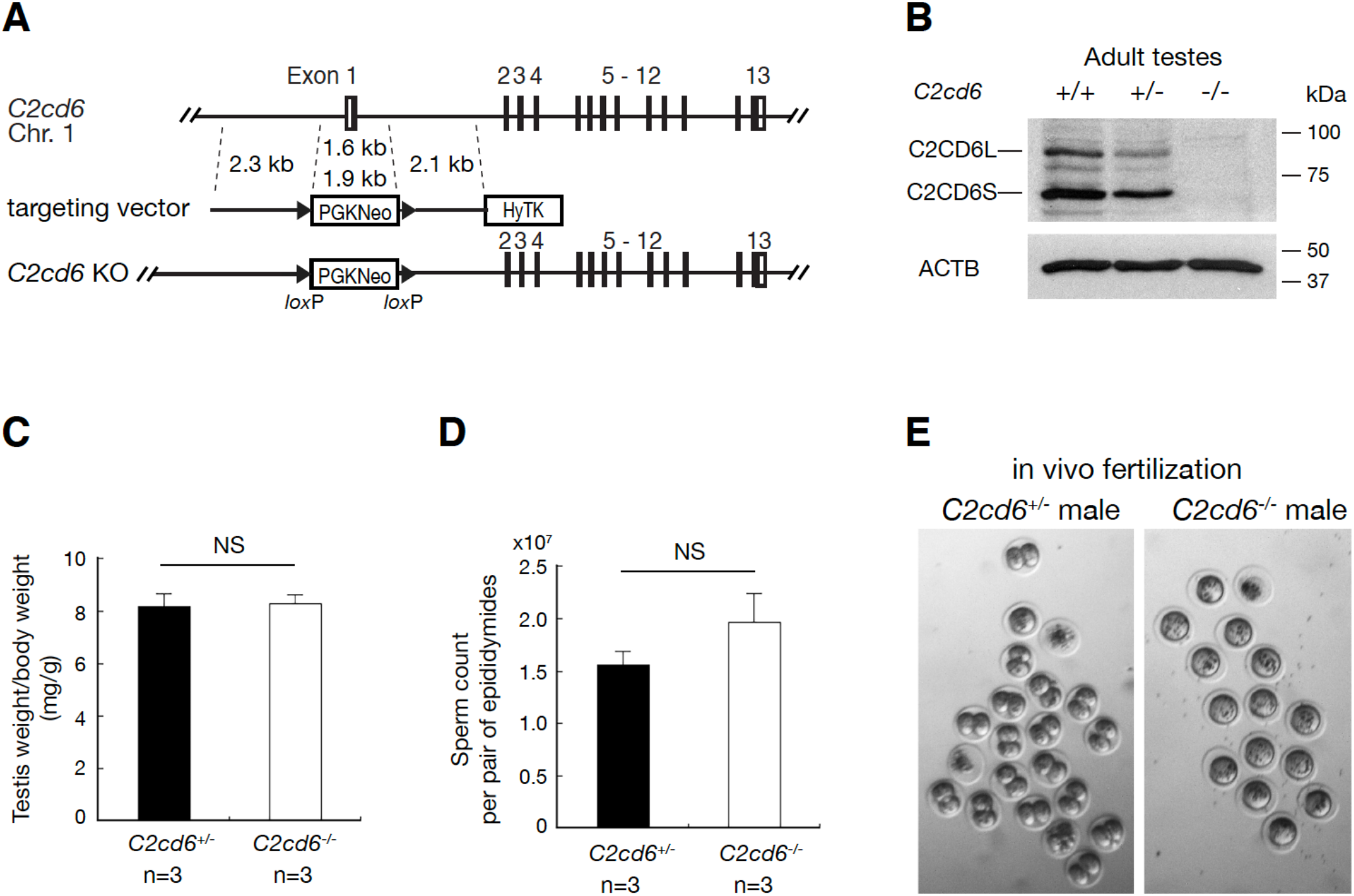
C2CD6 is essential for in vivo fertilization. (A) Targeted inactivation of the *C2cd6* gene. The 1.6-kb deleted region includes exon 1 (containing the initiating start codon) and 600-bp upstream of exon 1 (presumably the promoter region). The neomycin selection marker PGKNeo is flanked by *loxP* sites. HyTK provides negative selection by ganciclovir in ES cells. (B) Western blot analysis of C2CD6 in adult testes. ACTB serves as a loading control. (C) Testis weight normalized to bodyweight. (D) Sperm count. NS, not statistically significant. (I) In vivo fertilization assay. Embryos/unfertilized eggs were flushed from wild type females mated with either *C2cd6*^+/-^ or *C2cd6*^-/-^ males.

We generated polyclonal antibodies against a recombinant C2CD6 N-terminal 386-aa protein. Western blot analysis showed that C2CD6 was present in adult testis but not in ovary or somatic tissues in mice, demonstrating that C2CD6 is testis-specific (Fig. 1C). Interestingly, C2CD6 appeared as two major isoforms: C2CD6S (short, ∼60 kD) and C2CD6L (long, ∼85 kD). To determine the nature of these two C2CD6 isoforms, we performed RT-PCR, 3’RACE, and sequencing. The presence of two C2CD6 isoforms was due to alternative splicing (Fig. 1A). The *C2cd6s* transcript (GenBank accession number: NM_175200) consists of 13 exons and encodes a protein of 557 aa. The *C2cd6l* transcript (GenBank accession number: MW717645) is more complex and harbors alternative exon 13, alternative exon 1 of *Als2cr11b*, and splicing of an intron within *Als2cr11b* exon 5. As a result, the protein (691 aa) encoded by the *C2cd6l* transcript is larger than C2CD6S (557 aa) but smaller than ALS2CR11B (1729 aa). C2CD6S and C2CD6L share the first 533 aa and the predicted Ca^2+^-binding membrane targeting domain (Fig. 1B). Therefore, our C2CD6 antibody recognizes both isoforms. We then generated polyclonal antibodies against the C-terminal 200 aa of ALS2CR11B. Western blot analysis identified a protein band of ∼180 kD in testis but also in lung and brain at a low abundance, suggesting that *Als2cr11b* encodes a bona fide protein (Fig. 1C). Neither C2CD6 nor ALS2CR11B contain predicted transmembrane domains.

We immunostained sperm with anti-C2CD6 antibodies. Immunofluorescence analysis revealed that C2CD6 localized specifically to the principal piece of sperm flagellum (Fig. 1D), showing that C2CD6 is a novel component of sperm flagella. The C2CD6 signal is strong at the beginning of the principal piece (at the annulus region) and tappers off toward the end piece. The C2CD6 signal is specific, since it is absent in *C2cd6*-deficient sperm (Fig. 1D). Noticeably, two distinctive columns of C2CD6 can be appreciated in the principal piece by widefield fluorescence microscopy (Fig. 1D). These results demonstrate that C2CD6 is a new component of the sperm flagella.

### C2CD6 is essential for male fertility and in vivo fertilization

To study the functional requirement of *C2cd6*, we inactivated *C2cd6* by deleting a 1.6-Kb genomic region, including exon 1 (containing the initiating codon), through gene targeting in ES cells (Fig. 2A). Homozygous *C2cd6*^-/-^ mice were viable and grossly normal. While *C2cd6*^-/-^ females had normal fertility, *C2cd6*^-/-^ males were sterile. *C2cd6*^-/-^ males produced copulatory plugs, indicating normal mating behavior. Western blotting analysis showed that both C2CD6S and C2CD6L were absent in *C2cd6*^-/-^ testes, suggesting that the mutant is null (Fig. 2B). Testis weight and sperm count were comparable between *C2cd6*^+/-^ and *C2cd6*^-/-^ males (Fig. 2C and 2D). *C2cd6*-deficient sperm displayed normal morphology. Histology of *C2cd6*^-/-^ testes showed apparently normal spermatogenesis (data not shown) and thus C2CD6, unlike TEX11 (Yang et al., 2008), was not essential for meiosis.

*C2cd6*-deficient sperm were apparently motile, however, *C2cd6*^-/-^ males were sterile. To probe the cause of male infertility, we performed in vivo fertilization test. Wild type C57BL/6 females were injected with PMSG followed by hCG injection, mated with either *C2cd6*^+/-^ or *C2cd6*^-/-^ males (3 males per genotype), and copulatory plugs were checked. 24 hours after plug check, eggs/embryos were flushed from oviducts of plugged females. The number of two-cell embryos and one-cell embryos/eggs was counted. The majority of embryos (40/59 = 68%) from females plugged by *C2cd6*^+/-^ males were at the 2-cell stage, in contrast, only unfertilized eggs (a total of 51) were obtained from females plugged by *C2cd6*^-/-^ males (Fig. 2E). These data demonstrate that C2CD6 is required for fertilization *in vivo*.

To determine the capability of *C2cd6*-deficient sperm in egg activation and embryo development, we performed ICSI (intracytoplasmic sperm injection). Out of 50 oocytes injected with *C2cd6*-deficient sperm, 34 embryos reached the 2-cell stage and 20 of them further developed to the blastocyst stage. This result shows that *C2cd6*-deficient sperm can fertilize oocytes by ICSI and support embryo development.

### C2CD6 is required for in vitro fertilization and hyperactive motility

We next asked whether *C2cd6*-deficient sperm can fertilize oocytes in vitro. We performed in vitro fertilization (IVF) assays using cumulus-oocyte complexes from wild type CD1 females (Table 1). An average fertilization rate of 59 % vs 2 % was obtained when oocytes were incubated with *C2cd6*^+/-^ vs *Cdc26*-deficient sperm (Fig. 3A). Only embryos derived from *C2cd6*^+/-^ sperm developed to blastocysts (76%) after culture in KSOM media (Fig. 3B). The observed 2% fertilization rate obtained with *C2cd6*^-/-^ sperm was likely due to the low frequency of spontaneous parthenogenetic activation of oocytes (Xu et al., 1997). Therefore, these data indicate that C2CD6 is essential for fertilization in vitro.

**Table 1.** In vitro fertilization

| Genotype | Total oocytes | Unfertilized | Degraded | 2-cell embryos | Blastocysts |
| --- | --- | --- | --- | --- | --- |
| *C2cd6*<sup>+/-</sup> (n = 3) | 202 | 95 | 13 | 94 | 74 |
| *C2cd6*<sup>-/-</sup> (n =3) | 186 | 168 | 14 | 4 | 0 |

**Figure 3.**
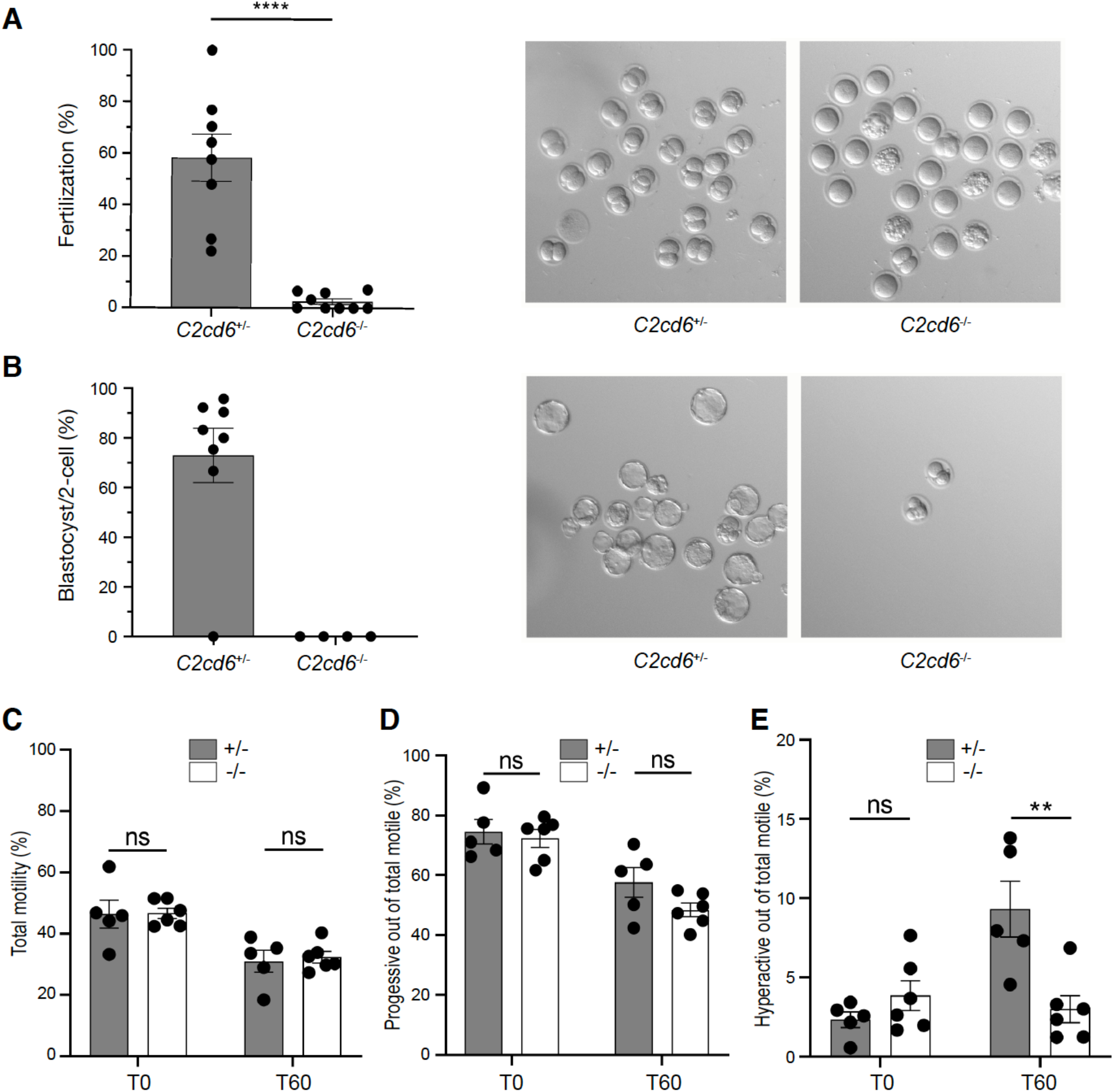
C2CD6 is required for in vitro fertilization and sperm hyperactivation. (A) In vitro fertilization. CD1 cumulus-oocyte complexes were incubated with *C2cd6*^+/-^ or *C2cd6*^-/-^ sperm. Fertilization rate is the percentage of oocytes inseminated that develop into 2-cell embryos after 24 hours of incubation. A representative image for each treatment is shown. ****p<0.0001. (B) The percentage of 2-cell embryos that develop to the blastocyst stage after culture in KSOM media. A representative image for each treatment is shown. (C) Percentage of motile sperm immediately after swim-out in TYH medium (T0) and after 60 minutes of incubation in capacitating conditions in TYH medium (T60). Sperm motility was analyzed by computer assisted sperm analysis (CASA). (D) Percentage of the motile sperm displaying progressive motility at T0 and at T60 of incubation in capacitating conditions. (E) Percentage of the motile sperm displaying hyperactive motility at T0 and at T60 of incubation in capacitating conditions. ns, not statistically significant, ** p<0.01.

To determine if the in vitro fertilization failure is related to sperm motility, we performed CASA (computer assisted sperm analysis) immediately after sperm swim-out from epididymis (T0) and after 60 minutes of incubation under capacitating conditions (T60). No differences in total or progressive sperm motility were found between *C2cd6*^+/-^ and *C2cd6*^-/-^ males (Fig. 3C and 3D).

Nevertheless, *C2cd6*-deficient sperm failed to acquire hyperactivated motility after 60 minutes of incubation under capacitating conditions (Fig. 3E). This could be the cause of the infertility phenotype in *C2cd6*^-/-^ males, since acquisition of hyperactivated motility is essential for fertilization.

We next evaluated the possibility of restoring fertility of the *C2cd6*-deficient sperm in vitro. We applied two sperm treatments prior to IVF that have been proven to restore fertility of other infertile and subfertile mouse models (Navarrete, Felipe A. et al., 2016;Navarrete, Felipe A. et al., 2019). Transient incubations with Ca^2+^ ionophore (ionophore) and sperm energy restriction (SER) produced a moderate increase in the fertilization rates of *C2cd6*-deficient sperm: 6.3% and 18.8% respectively (Table 2). Strikingly, the combination of SER and ionophore treatments of *C2cd6*-deficient sperm induced an average fertilization rate of 58.8% as a total of 50 2-cell embryos out of 85 inseminated oocytes were obtained (Table 2). To analyze embryo development in vitro, the obtained 2-cell embryos were further cultured in KSOM media. We observed a blastocyst development rate of 22% in the ionophore treatment-derived embryos, 100% in the SER-derived embryos, and 63% in the SER + ionophore-derived embryos. Although each sperm treatment was able to overcome the infertility phenotype of *C2cd6*^-/-^ males in vitro, the combination of SER and ionophore treatments was the most effective.

**Table 2.** Ionophore and SER treatments

|  | Genotype | Control | Control +<br>Ionophore | SER | SER +<br>Ionophore |
| --- | --- | --- | --- | --- | --- |
| Fertilization<br>(%) | *C2cd6<sup>+/-</sup>* | 222/405 (54.8) | 46/48 (95.8) | - | - |
|  | *C2cd6<sup>-/-</sup>* | 10/389 (2.6) | 6/96 (6.3) | 15/80 (18.8) | 50/85 (58.8) |
| Blastocyst/2-<br>cell (%) | *C2cd6<sup>+/-</sup>* | 144/222 (64.9) | 43/46 (93.5) | - | - |
|  | *C2cd6<sup>-/-</sup>* | 0/10 (0) | 2/5 (22.2) | 14/14 (100) | 31/50 (62.7) |

### C2CD6-dependent CatSper assembly in sperm flagella

CatSper, the flagellar Ca^2+^ ion channel, localizes to the principal piece (Fig. 4A) (Chung et al., 2017;Ren et al., 2001). C2CD6 localization to the sperm flagella (Fig. 1D) is strikingly similar to CatSper localization. Moreover, using comparative proteomics, C2CD6 was shown to be one of the proteins displaying reduced abundance in CatSper1-deficient sperm (Hwang et al., 2019). Therefore, we examined CatSper1 localization in the absence of C2CD6 by immunofluorescence. The CatSper1 signal was severely reduced in *C2cd6*^-/-^ sperm (Fig. 4A). We also performed super-resolution imaging analysis of CatSper1 and C2CD6 localization in sperm flagellum. As previously reported, CatSper1 formed quadrilateral columns in wild type flagellum (Fig. 4B). C2CD6 also appeared in columns but was less organized than CatSper1 columns in wild type (Fig. 4B). In *C2cd6*^-/-^ sperm, CatSper1 signals were sharply reduced, disorganized, discontinuous, and preferentially distributed to the distal end of the principal piece (Fig. 4B).

**Figure 4.**
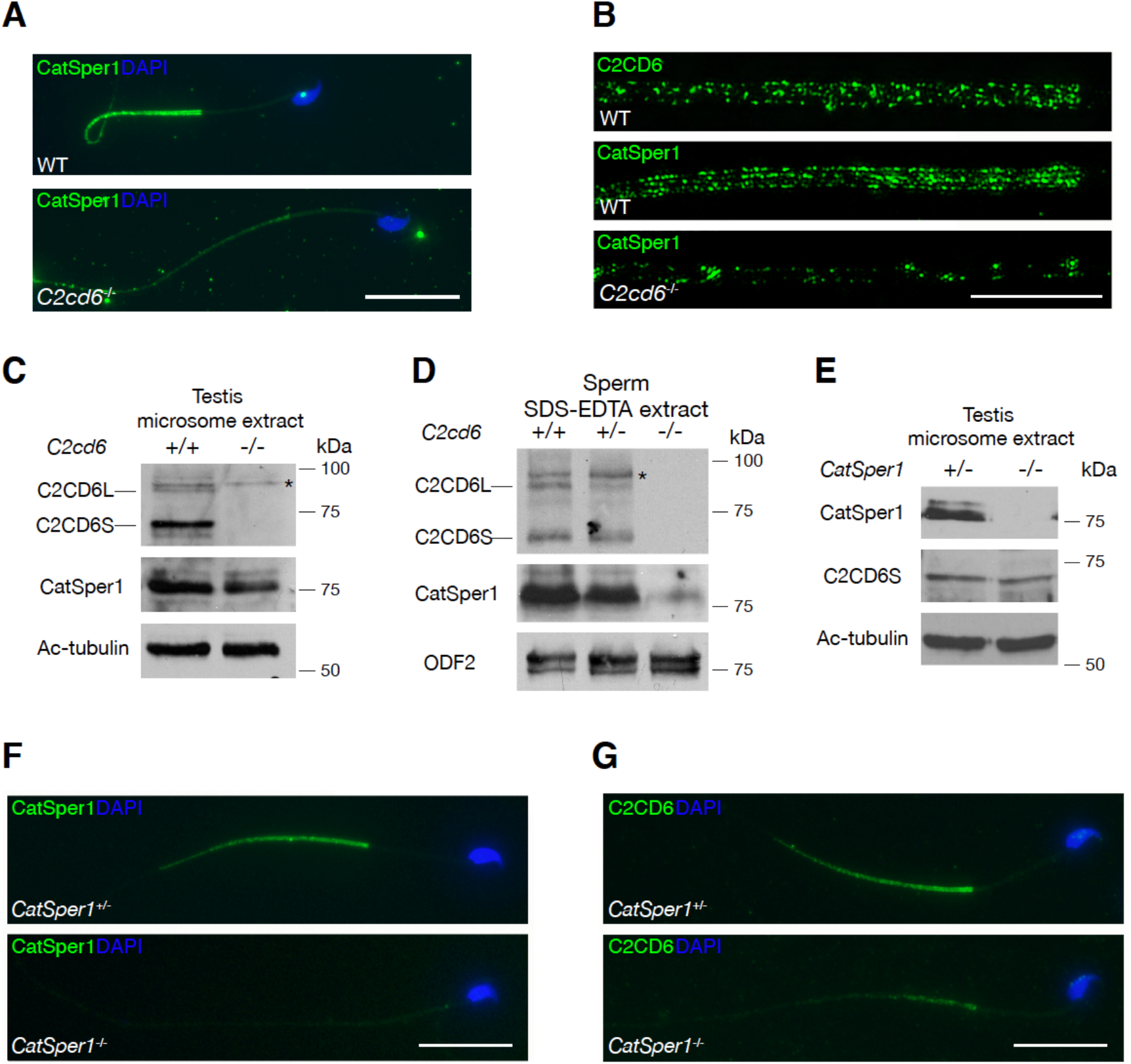
C2CD6 is required for CatSper assembly in sperm flagella. (A) Immunofluorescence analysis of CatSper1 in wild type and *C2cd6*-deficient sperm. Scale bar, 25 μm. (B) Super-resolution localization of C2CD6 and CatSper1 in sperm. Scale bar, 5 μm. (C) Western blot analysis of C2CD6 and CatSper1 in testis microsome fractions. Acetylated tubulin serves as a loading control. (D) Western blot analysis of C2CD6 and CatSper1 in sperm extracts. ODF2, a component of flagellar outer dense fibers, serves as a loading control (Cao et al., 2006). (E) Immunofluorescence analysis of CatSper1 in *CatSper1*^+/-^ and *CatSper1*^-/-^ sperm. (F) Localization of C2CD6 in *CatSper1*^+/-^ and *CatSper1*^-/-^ sperm. Scale bars, 25 μm.

CatSper complex is partitioned into microsomes (vesicle-like structures formed from pieces of endoplasmic reticulum) in testis extracts (Chung et al., 2017). We prepared microsomes from wild type and *C2cd6*^-/-^ testes (Fig. 4C). C2CD6, like CatSper1, was present in testicular microsome fractions. In addition, CatSper1 was abundant in microsome fractions from *C2cd6*^-/-^ testes (Fig. 4C), suggesting that synthesis of CatSper is not affected in *C2cd6*^-/-^ testes. We next performed Western blotting analysis of sperm extracts. As expected, both C2CD6 and CatSper1 were present in wild type and *C2cd6*^+/-^ sperm extracts (Fig. 4D). However, CatSper1 abundance was dramatically reduced in *C2cd6*^-/-^ sperm extract (Fig. 4D). Taken together, our results demonstrate that the CatSper ion channel complex is abundant in the testis but fail to incorporate into sperm flagella in the absence of C2CD6. Therefore, C2CD6 is required for assembly of the CatSper channel complex in sperm flagella.

We next sought to address whether CatSper is required for C2CD6 localization. Both CatSper1 and C2CD6 were detected in *CatSper1*^+/-^ testis microsome extract (Fig. 4E). C2CD6 was present in *CatSper1*^-/-^ testis microsome extract and its abundance was comparable with that in *CatSper1*^+/-^ testis (Fig. 4E). As expected, CatSper1 was absent in *CatSper1*^-/-^ flagellum (Fig. 4F). C2CD6 was sharply reduced in *CatSper1*^-/-^ flagellum (Fig. 4G). This result is consistent with the reduced abundance of C2CD6 (formerly known as ALS2CR11) in *CatSper1*^-/-^ sperm shown by quantitative proteomic analysis (Hwang et al., 2019). These results demonstrate that CatSper is critical for C2CD6 localization in sperm flagella. Therefore, the inter-dependent localization of C2CD6 and CatSper1 suggests that C2CD6 might be an essential component of the CatSper complex.

### C2CD6 interacts with components of the CatSper complex

To investigate the connection of C2CD6 with the CatSper channel complex, we co-expressed C2CD6 with CatSper complex components in HEK293T cells and tested their interaction by co-immunoprecipitation. C2CD6 and all CatSper components are only expressed in testis and remain in sperm but are absent in somatic cells such as 293T cells. The abundance of C2CD6 in transfected HEK293T cells was low but can be dramatically enriched by immunoprecipitation (Fig. 5). The full-length C2CD6 migrated at 75 kDa and a slightly smaller isoform was also present in the immunoprecipitated fraction (Fig. 5A). CatSper1 was present in the C2CD6 complex (Fig. 5A). CatSper2 was also in complex with C2CD6 but the association was weak (Fig. 5B). CatSper3 was strongly associated with C2CD6 (Fig. 5C). CatSper4 was readily detected in C2CD6-immunoprecipated proteins, indicating strong interaction between CatSper4 and C2CD6 (Fig. 5D). EFCAB9 was abundant in C2CD6-immunoprecipated proteins, showing that EFCAB9 is strongly associated with C2CD6 (Fig. 5E). However, we did not detect interaction between C2CD6 and CatSperz (TEX40) (Fig. 5F). Collectively, C2CD6 interacts with the core components of the CatSper complex (Fig. 5G).

**Figure 5.**
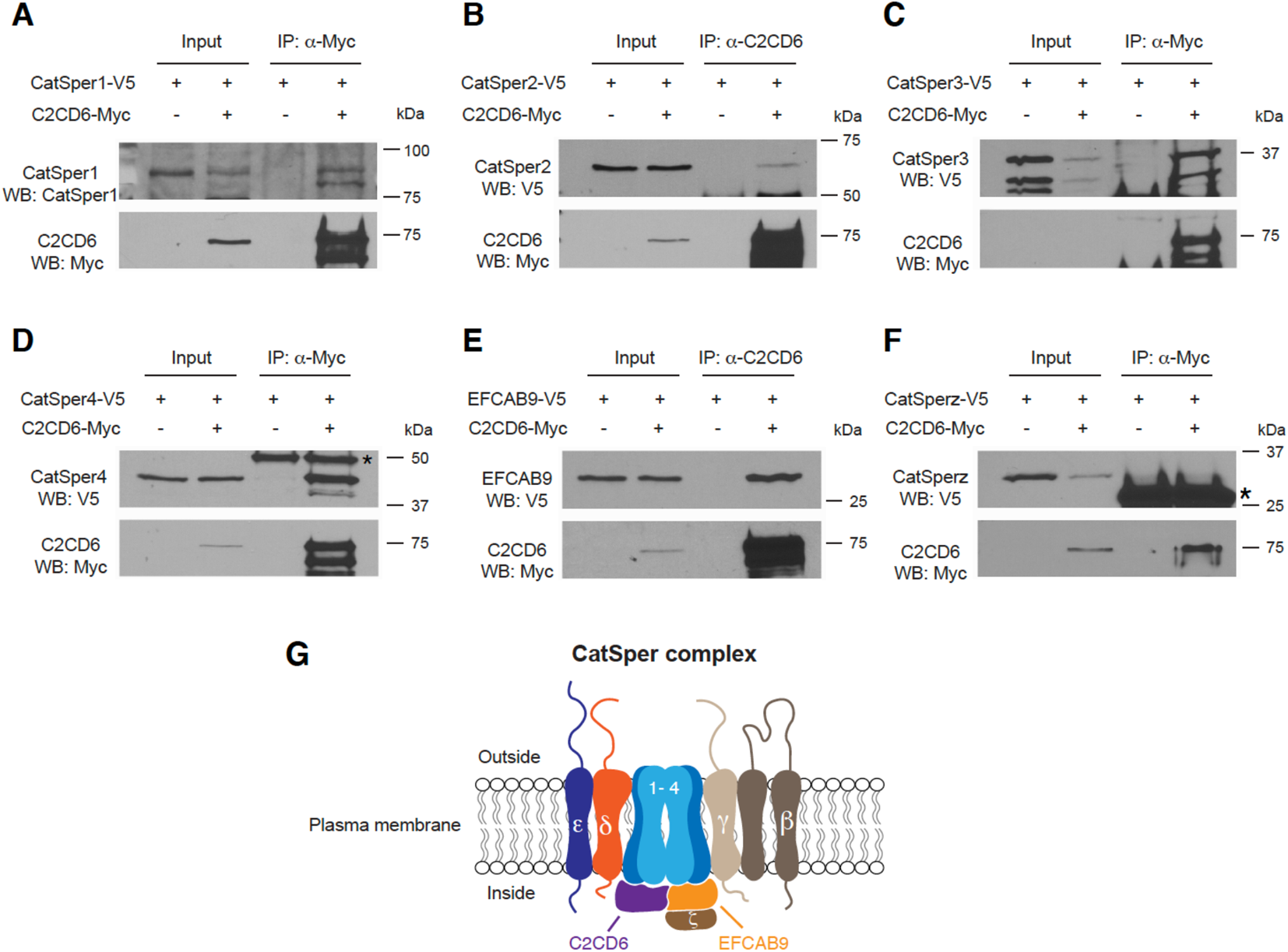
C2CD6 interacts with subunits of the CatSper channel complex. HEK293T cells were transfected with indicated expression constructs and protein extracts were used for immunoprecipitation. Coimmunoprecipitation of C2CD6 with CatSper1 (A), CatSper2 (B), CatSper3 (C), CatSper4 (D), and EFCAB9 (E). Asterisk in panel D indicates Ig heavy chain. (F) C2CD6 is not associated with CatSperz. Asterisk indicates Ig light chain. (G) A working model of the CatSper channel complex. Eleven known subunits of the CatSper channel including C2CD6 are shown. C2CD6 interacts with CatSper 1-4 and EFCAB9.

## Discussion

Here we demonstrate that C2CD6 is a novel and essential subunit of the CatSper complex, in addition to the ten known subunits (Fig. 5G). Four core α subunits, CatSper1-4, form the membrane-spanning Ca^2+^-selective pore. C2CD6 interacts with all four α subunits suggesting that it directly binds to the CatSper channel core. In addition, C2CD6 interacts with EFCAB9 but not with CatSperZ, a partner of EFCAB9. We postulate that C2CD6 plays several possible roles in the CatSper channel function. First, C2CD6 might be a critical structural subunit of CatSper. The decrease of CatSper1 in the C2CD6-deficient sperm and the decrease of C2CD6 protein in CatSper1-deficient sperm suggest that without C2CD6, the CatSper complex is not fully assembled. CatSper subunits localize in linear quadrilateral domains along the sperm principal piece, and the localization of CatSper1 is disrupted in animals lacking other subunits such as EFCAB9, CatSper?, and CatSperε (Chung et al., 2011;Chung et al., 2014;Chung et al., 2017). Consistent with C2CD6 as a novel subunit of the CatSper channel complex, we find that Catsper1 is drastically reduced in *C2cd6*-deficient sperm and displays disrupted quadrilateral domain localization in the principal piece. Second, C2CD6 contains a calcium-dependent membrane-targeting C2 domain. This domain is found in signaling proteins that interact with the cellular membrane (Nalefski and Falke, 1996). This raises the possibility that C2CD6 might facilitate targeting of assembled CatSper complexes to the sperm flagellar plasma membrane or insertion into the membrane. Indeed, CatSper1 is present in testis but not in sperm from *C2cd6*^-/-^ males, indicating that the CatSper complex is unstable without C2CD6. Similar results were found when each of the other CatSper subunits was individually removed. In each of these cases, the remaining subunits are expressed in the testes but are absent in mature sperm (Qi et al., 2007). Third, C2CD6 might function as a Ca^2+^ sensor for CatSper independently or as a complex with EFCAB9. Notably, the CatSper channel is compromised but still conducts currents in EFCAB9-deficient sperm (Hwang et al., 2019). In EFCAB9-deficient sperm, C2CD6 might be responsible for Ca^2+^ sensing.

C2CD6 exists as at least two isoforms. Both isoforms contain the C2 domain and are present in sperm. It is not known whether these two isoforms are functionally redundant or have isoform-specific functions. Intriguingly, CatSperδ also has two isoforms resulting from alternative splicing (Chung et al., 2011). In addition, C2CD6 (previously known as ALS2CR11) is tyrosine phosphorylated upon capacitation (Chung et al., 2014). The physiological consequence of this phosphorylation on C2CD6 is unknown. The *Als2cr11b* gene encodes a bona fide protein in testis (Fig. 1C). Like *C2cd6, Als2cr11b* is conserved in vertebrates. Future study is necessary to investigate whether ALS2CR11B is a subunit of the CatSper complex. Genetic ablation of *Als2cr11b* alone without disruption of *C2cd6* is challenging, because *Als2cr11b* shares exons with *C2cd6l* (Fig. 1A).

*C2cd6*^-/-^ sperm do not fertilize oocytes in vivo or in vitro. The sterility phenotype is caused by the failure in induction of hyperactivated motility during sperm capacitation. Hyperactivated motility is characterized by a high-amplitude asymmetrical beating of the sperm flagellum (Suarez and Osman, 1987). This flagellar beating pattern is mainly regulated by sperm intracellular Ca^2+^ concentrations ([Ca^2+^]_i_), which are maintained by ion channels and pumps in the plasma membrane (Visconti, Pablo E. et al., 2011). Two of the most important sperm [Ca^2+^]_i_ regulators are the CatSper channel that is essential for the entrance of Ca^2+^ to the sperm (Ren et al., 2001), and the Ca^2+^ efflux pump PMCA4 that is required for Ca^2+^ clearance (Wennemuth et al., 2003). The proper assembly and function of the CatSper channel are essential for acquisition of sperm hyperactivated motility and fertility (Chung et al., 2011;Chung et al., 2017;Hwang et al., 2019;Ren et al., 2001). Consistent with our conclusion that C2CD6 is a novel subunit of the CatSper channel complex, *C2cd6*-deficient sperm fail to achieve hyperactivated motility after incubation under capacitating conditions.

Ca^2+^ is a second messenger with pivotal roles in activating (or inhibiting) downstream effectors during sperm capacitation (Navarrete, F. A. et al., 2015). The transient treatment of sperm with Ca^2+^ ionophore A23187 can bypass the activation of the main molecular pathway critical for acquisition of fertilizing capacity during sperm capacitation: the cAMP/PKA pathway (Tateno et al., 2013). This transient sperm ionophore treatment was applied prior to IVF and successfully reversed the male sterility phenotype of *CatSper1*^-/-^, *sAC*^-/-^, and *Slo3*^-/-^ mice as moderate fertilization was achieved (Navarrete et al., 2016). In line with these previous observations, the ionophore treatment prior to IVF in *C2cd6*-deficient sperm reversed the sterility phenotype, indicating that an increase of intracellular Ca^2+^ is sufficient to restore fertility in this mouse model. We have recently developed another sperm treatment that improves fertilization rates and embryo development of sub-fertile mouse models by manipulation of the sperm metabolism (Navarrete et al., 2019). When applied prior to IVF in *CatSper1*^-/-^ sperm, this Sperm Energy restriction and Recovery (SER) treatment was not able to restore fertility; however, a combination of the ionophore A23187 and the SER treatments induced a synergistic effect on fertilization rates and embryo development in the *CatSper1*^-/-^ model (Navarrete et al., 2019). The same synergistic effect was observed on *C2cd6*-deficient sperm. Interestingly, while the SER treatment did not rescue the sterility phenotype of *CatSper1*^-/-^ mice (Navarrete et al 2019), the application of SER treatment alone was able to overcome the sterility phenotype of the *C2cd6*^-/-^ mice. This could be related to our recent findings that SER treatment induces an elevation of [Ca^2+^]_i._ in mouse sperm from wild type and *Catsper1*^-/-^ animals (Sánchez-Cárdenas et al., 2021).

The CatSper channel is essential for sperm hyperactivation and male fertility in both mice and humans. The CatSper subunits are only expressed in testis and sperm. Traditionally, ion channels are druggable targets. For these reasons, the CatSper channel has been proposed as a target for male contraception with minimal side effects. However, reconstitution of CatSper in a heterologous system has not been achieved, despite that CatSper was discovered two decades ago (Ren et al., 2001). The lack of a heterologous system impedes drug discovery efforts for small molecule inhibitors of CatSper. The challenges for developing a CatSper heterologous system are several fold. First, the CatSper ion channel is extremely complex. It is still possible that not all CatSper-associated proteins are known. C2CD6 is the newest CatSper subunit. Second, the CatSper assembly might require chaperones. The CatSper complex is associated with a testis-specific chaperone – HSPA2 (Chung et al., 2011;Zhu et al., 1997). Third, sperm flagellum is a unique ciliary structure. CatSper forms organized linear domains along the principal piece. Organization of these nanodomains might depend on other flagellar unique structures such as fibrous sheath, which are absent in heterologous cells. The identification of C2CD6 might facilitate successful development of a heterologous CatSper system, which is not only critical for drug development but also provide an amenable system to dissect the mechanistic role of each subunit in CatSper.

## Acknowledgements

We thank Dejian Ren for anti-CatSper1 antibody. This study was supported by NIH/National Institute of General Medical Sciences GM118052 (PJW), Eunice Kennedy Shriver National Institute of Child Health and Human Development HD069592 and HD068157 (PJW), HD38082 and HD088571 (PEV), and National Research Service Award T32 GM108556 (DAT).

## Author contributions

F.Y. and P.J.W. conceptualized the study. F.Y. generated and characterized the *C2cd6* knockout mice. M.G.G., D.A.T., and P.E.V. contributed the CASA, IVF, and in vitro sperm treatment data; N.A.L. performed blastocyst injection and generated chimeric mice; G.R. contributed to the super resolution microscopy experiments; P.J.W., F.Y., M.G.G., and P.E.V. wrote the manuscript. All the authors commented on the manuscript.

